# Temporal Requirement for Stearoyl-CoA Desaturase 1 in Oligodendrocyte Development but Not Myelin Maintenance

**DOI:** 10.1101/2025.09.18.676928

**Authors:** Iris Farnum, Thaddeus J. Kunkel, Manideep Chavali

## Abstract

Stearoyl-CoA desaturase 1 is a rate-limiting enzyme in monounsaturated fatty acid synthesis, which is crucial for membrane biosynthesis. Here we show an early requirement for Scd1 in oligodendroglial cells during developmental myelination. Using oligodendrocyte progenitor cell (OPC) specific conditional knockout model of *Scd1*, we observed a myelination delay during CNS development. Genetic ablation of OPC-specific *Scd1* resulted in oligodendrocyte maturation delay and hypomyelination within forebrain white matter tracts and optic nerve. Interestingly, although expressed at high levels within the mature oligodendrocytes, Scd1 was dispensable in maintenance of oligodendrocytes and axonal myelination, as loss of mature oligodendrocyte specific Scd1 showed no effect on myelin maintenance or oligodendrocyte survival. Together, our results suggest that Scd1 function is temporally restricted to the developmental period when oligodendrocytes undergo differentiation and active myelination but becomes dispensable for maintaining established myelin.

## 1. Introduction

During central nervous system development, oligodendrocyte precursor cells undergo morphological changes as they differentiate into mature oligodendrocytes, extending processes that can myelinate several axonal segments simultaneously. This process requires massive myelin membrane synthesis. Myelination by oligodendrocytes involves formation of lipid-rich membrane sheaths that wrap around axons. This compact myelin wrapping enables rapid saltatory conduction and also allows efficient metabolite transport to the underlying axons. Myelin membrane is unique among other cellular membranes, comprising nearly 70% lipids including cholesterol, phospholipids with specific fatty acid composition that regulate membrane stability and fluidity for appropriate axonal wrapping (Chrast et al., 2011; Gielen et al., 2006; O’Brien & Sampson, 1965; Stadelmann et al., 2019; Williams & Deber, 1993).

The biophysical properties of the myelin membrane depend critically on the lipid composition (Bosio et al., 1998; Nottar Escobar et al., 2025). During the early myelination phase, oligodendrocyte membrane must maintain sufficient fluidity to extend across axons of various calibers and then wrap around them (Montani, 2021; Sherman & Brophy, 2005; Young et al., 2013). Next, these membrane wraps become highly stable to achieve a compacted structure of mature myelin. Monounsaturated fatty acids (MUFAs) are known to provide this essential cell membrane fluidity (Scott et al., 2022; Zhao et al., 2016). Stearoyl-CoA desaturases (Scd1-4) are rate-limiting enzymes that catalyze the synthesis of these MUFAs (Ntambi, 1995, 1999). Among these enzymes, Scd1 and Scd2 show high expression within the CNS cell types, particularly within the oligodendrocyte lineage cells and astrocytes (Cahoy et al., 2008; Naffaa et al., 2022).

The importance of lipid metabolism in oligodendrocyte function is well established(Camargo et al., 2017; Montani, 2021; O’Brien & Sampson, 1965; Zhou et al., 2020). Prior work has shown that disruption of cholesterol and lipid synthesis within oligodendrocyte lineage and astrocytes prevents normal myelination (Camargo et al., 2017). Targeted disruption in fatty acid synthesis pathways is also known to cause hypomyelination, structural defects in myelin wrapping and oligodendrocyte death (Camargo et al., 2017; Dimas et al., 2019). However, despite high expression of Scd1 and Scd2 in oligodendrocytes, their specific contributions to these processes remain unexplored. Oligodendrocytes have vastly different metabolic requirements during active developmental myelination versus the maintenance of established myelin sheaths in adulthood (Funfschilling et al., 2012; Harris & Attwell, 2012; Rinholm et al., 2011; Saher et al., 2005). During development, they rapidly synthesize large quantities of membrane lipids. In contrast, mature oligodendrocytes primarily maintain existing myelin with presumably lower biosynthetic demands (Ando et al., 2003; Chrast et al., 2011; Snaidero et al., 2014; Young et al., 2013). Whether the same metabolic pathways support both phases or whether stage-specific programs exist remains unknown.

Here we used conditional knockout approaches to understand the temporal requirements for Scd1 in oligodendrocyte development and myelin maintenance. By targeting Scd1 deletion to oligodendrocyte precursor cells during early development or to mature oligodendrocytes after developmental myelination, we uncovered a temporal specificity of Scd1 function. We found that Scd1 contributes to efficient OPC differentiation and developmental myelination, with its loss causing a reduction in mature oligodendrocyte numbers and myelin thickness. In contrast, Scd1 becomes dispensable for myelin maintenance. These findings reveal temporal differences in metabolic requirements between developing and mature oligodendroglial cells.

## 2. Materials and Methods

### Animals

All animal procedures were performed according to the Oregon Health and Science University guidelines under Institutional Animal Care and Use Committee (IACUC) approved protocol. Animals of both sexes were analyzed. Following transgenic lines were employed in this study: *Scd1 floxed* mouse line was previously described in Miyazaki et al., 2007 and generously donated by Dr. James Ntambi’s Lab. *PDGFRa-Cre*^*ER*^ mouse line was described in Kang et al., 2010 and obtained from Jackson Laboratory (strain # 018280). *Plp1-Cre*^*ER*^ mouse line was previously described in Doerflinger et al., 2003 and obtained from Jackson Laboratory (strain # 05975). *R26R-EYFP* mouse line was described in Srinivas et al., 2001 and obtained from Jackson Laboratory (strain # 006148). Cre-mediated deletion of *Scd1* was induced by intraperitoneal injection of tamoxifen (Sigma #T5648) dissolved in corn oil (Sigma #C8267) and analyzed according to the timelines shown in the figures. For induction in *OPC-Scd1 cKO* mice, tamoxifen was administered at 100µg/day from P1–P3. For induction in *OL-Scd1 cKO mice*, tamoxifen was administered at 1mg/day from P13–P15 or from P30–P35.

### Immunohistochemistry

For immunohistochemical analysis on brain sections, transcardiac perfusion was performed on anesthetized mice with 4°C PBS, followed by 4% paraformaldehyde. Brains were isolated and post-fixed overnight in 4% paraformaldehyde and cryo-protected in 30% sucrose for atleast 24 hours prior to OCT embedding. Coronal brain sections were obtained at 25μm thickness. Antigen retrieval on the tissue sections was performed for 2 minutes at 95°C in 10mM Sodium Citrate buffer (pH 6.0) in a Pelco-Biowave (TedPella). Tissue sections were washed in 4°C PBS and then blocked for 1 hour in 5% horse serum buffer and 0.3% Triton X-100 in PBS. Next, sections were incubated with primary antibodies overnight at 4°C in fresh blocking solution. Sections were washed in PBS containing 0.3% Triton X-100 three times for five minutes each, followed by 1 hour incubation in species specific-secondary antibodies (1:500 Invitrogen). Sections were then washed again five times and were mounted using DAPI-FluoromountG (Southern Biotech). Primary antibodies used are anti-Olig2 (1:100; R&D Systems, AF2418), anti-CC1 (1:500; Millipore, OP80), anti-BCAS1 (1:500; Santa Cruz Biotechnology, sc-393740), anti-MBP (Millipore, MAB386), and anti-NF (1: 2000; EnCor Biotechnology, CPCA-NF-L).

### RNAscope

Fixed frozen mouse brain sections were analyzed by RNAscope Multiplex Fluorescent Kit V2 (Advanced Cell Diagnostics, 320850), with Pdgfrα-Mm probe (Advanced Cell Diagnostics, 480661-C3), Plp1-Mm probe (Advanced Cell Diagnostics, 428181-C1), and Scd1-Mm probe (Advanced Cell Diagnostics, 461641-C2). Briefly, cryosections were mounted on glass slides and baked at 60°C for 30 mins and were further fixed in 4% paraformaldehyde at 4°C for 15 minutes. Next, serial dehydration was performed in 50%, 70% and 100% ethanol for 5 minutes each at room temperature, followed by air drying. Next, target retrieval was performed with RNAscope reagents for 2 minutes at 95 °C. After a quick wash in the PBS, endogenous peroxides were blocked with a H_2_O_2_ treatment. Slides were washed twice in RNase free water before Protease-III (ACDBio) treatment for 10 mins at 40°C. Probes listed above were diluted at 1:50 ratio in a channel 1 probe. Hybridization with the probe mix was performed in HybEZ oven for 2 hours at 40°C. Next, slides were washed for 2 mins in wash buffer (ACDBio) twice before proceeding to the amplification steps according to manufactures protocol. Fluorescent detection was performed with Opal fluorophores (Akoya Biosciences) at a 1:500 dilution. Slides were then washed two times before proceeding with IHC labelling as described above.

### Electron Microscopy and Analysis

For electron microscopy analysis, animals were perfused with 20ml of 0.1 M PBS, pH 7.4 and next by 30 mL of freshly prepared cold fixative solution composed of 4% paraformaldehyde in 0.1M PBS, pH 7.4. Optic nerves were dissected and placed into fixative solution containing 2% paraformaldehyde and 2% glutaraldehyde for 24hr. The optic nerves were then placed in a long-term fixative solution with 1.5% paraformaldehyde, 1.5% glutaraldehyde, 1.5% sucrose, and 1.5% calcium chloride in 0.1M cacodylate buffer and water until resin embedded. Ultrathin sections were analyzed on Tecnai-T12 transmission electron microscope. For each myelinated axon, the g-ratio was calculated by dividing the axonal diameter by the total myelin fiber diameter (defined by the outer limit of the myelin sheath). Each group consisted of at least 3-4 mice/group and atleast 500 axons were analyzed (see figure legend for details).

### Microscopy and Cell Counting

Microscopy images were obtained on either a Lecia SP8 confocal microscope or Zeiss Axio Imager M2 widefield microscope. Images obtained were analyzed on FIJI (Image J). All images were obtained with same exposure levels. Brightness or contrast modifications on the images which evenly were applied between all control and experimental groups. Rostro-caudal level matched brain sections were used for imaging and analysis. For representative images a cartoon with highlighted region from which the images were obtained are shown in the insets. Atleast 3-6 fields from atleast 4-6 sections were analyzed per animal. All the cell counts presented are the numbers cells quantified from a field of 0.6mm^2^ area or percentage of cells within the indicated population. All analysis was performed on the FIJI software.

### RNA Sequencing

For RNA sequencing, P10 mice were transcardially perfused with PBS and optic nerves were dissected. Total RNA was isolated using TRIzol (Invitrogen) and purified with RNeasy Kit (Qiagen). Library preparation and sequencing were performed by Novogene using NovaSeq X Plus with 150bp paired-end reads, generating 41-64 million raw reads per sample. Read quality was assessed using FastQC (Andrews S, 2010). Reads were aligned to the mouse reference genome (GRCm39) using HISAT2 (Kim et al., 2015) with default parameters. Gene counts were generated using FeatureCounts (Liao et al., 2014). Differential expression analysis was performed using DESeq2, with genes considered significant at FDR<0.05. Gene ontology analysis was performed using PANTHER (Ashburner et al., 2000; Thomas et al., 2022), and gene set enrichment analysis (GSEA) (Subramanian et al., 2005) was conducted using pre-ranked gene lists from the DESeq2 differential expression statistics.

### Fatty Acid Analysis

For Fatty acid analysis, P10 mice were transcardially perfused with PBS and dissected optic nerve and corpus callosum tissue isolates were pooled together. Snap frozen samples were analyzed by Creative Proteomics Facility (New York). Briefly, samples were transferred glass vials containing Tritricosanoin as an internal standard (tri-C23:0 TG). These samples are then homogenized, and fatty acids were extracted following the modified Folch method. A portion of the organic layer was transferred to screw-cap glass vials and dried in a speed vac. After samples were dried, a mixture of methanol containing 14% boron trifluoride, toluene, methanol at 35:30:35 v/v/v (Sigma-Aldrich) was added. The vial was briefly vortexed and heated to 100°C for 45 minutes. Samples were allowed cool, and mixture of hexane (EMD Chemicals) and HPLC grade water was added to the tubes, followed by vertexing and centrifugation to help layer separation. An aliquot of the hexane layer was transferred to a gas chromatography vial. Fatty acids were identified by comparison with a standard mixture of fatty acids in the samples which was added prior the extraction (GLC OQ-A, NuCheck Prep) which was also used to determine individual fatty acid calibration curves.

### Statistics

All the data are represented as mean ± standard error of the mean (S.E.M.), and the number of animals per genotype is indicated in figure legends. Statistical analysis was performed using the two tailed unpaired Student’s t-test and data plotting was performed using GraphPad Prism 8.0 software. In all figure legends, p values are indicated by *p<0.05, ****p<0.0001.

## 3. Results

### 3.1 Loss of OPC-Specific Scd1 Results in Reduced OL Maturation During Early Development

To examine the role of Scd1 in oligodendrocyte development, we generated an OPC-specific conditional knockout mouse line (*OPC-Scd1 cKO*) using the tamoxifen-inducible *Pdgfrα-Cre*^*ER*^ line to delete Scd1 in oligodendrocyte precursor cells (OPCs). Tamoxifen was administered from P1-P3 to induce Cre-mediated recombination during the early postnatal period when OPCs are actively proliferating and beginning to differentiate (**Figure 1A**). Expression analysis showed that Scd1 transcript is expressed within *Pdgfrα*+ OPCs, BCAS1+ pre-myelinating oligodendrocytes **(Figure 1B)** and also within the *Plp1*-expressing mature oligodendrocytes in the control mice (**Supplementary Figure 1B**). These transcripts were reduced in the oligodendrocyte lineage cells of *OPC-Scd1 cKO* mice (**Figure 1B and Supplementary Figure 1A**).

**Figure 1.**
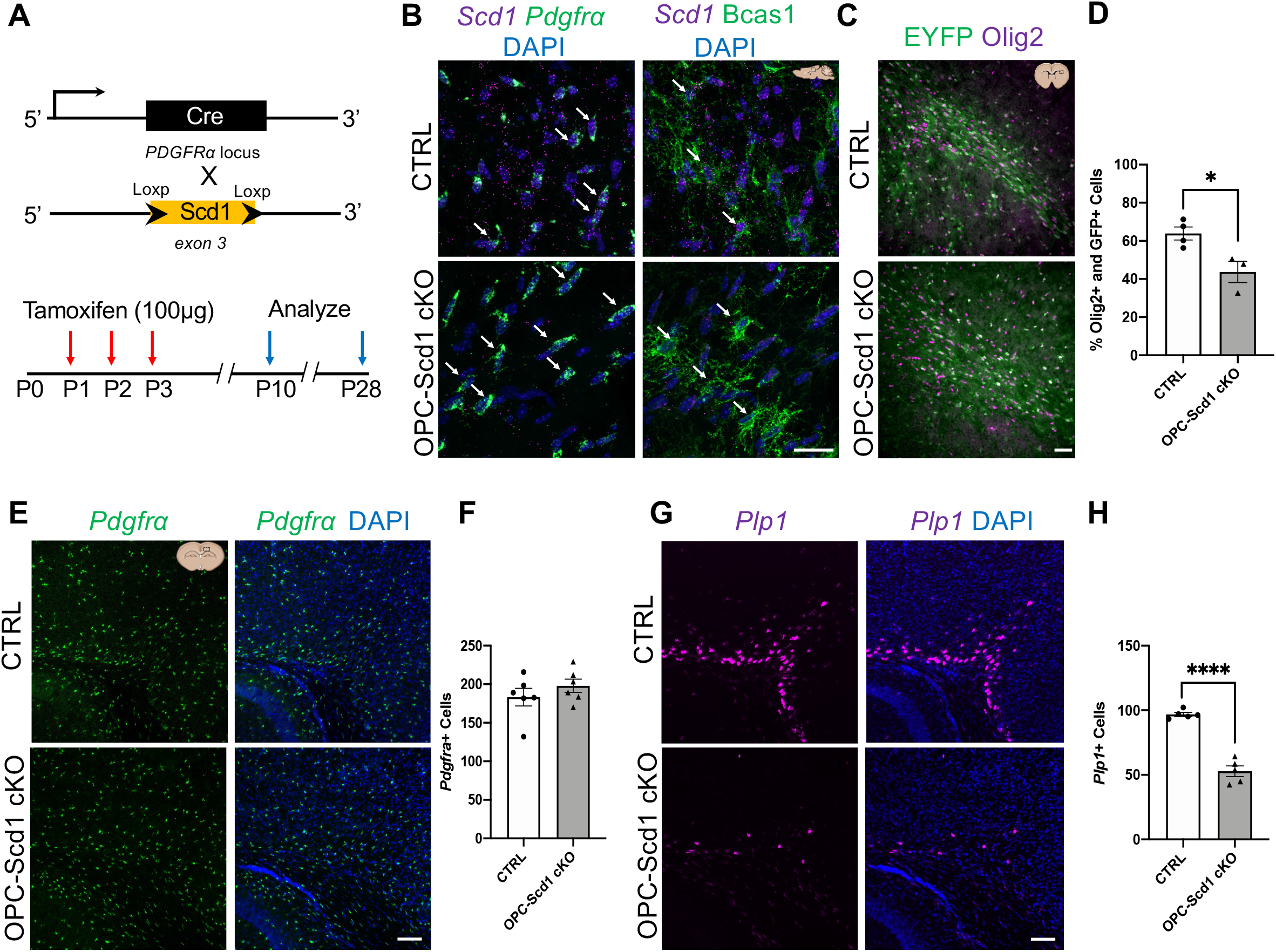
OPC-specific Scd1 deletion results in reduced oligodendrocyte maturation during early postnatal development. (A) Experimental design showing Pdgfrα-Cre^ER^-mediated genetic ablation of Scd1 (targeting exon 3) (referred to as OL-Scd1 cKO). Tamoxifen administration is performed from P1-P3. These animals are referred to as OPC-Scd1 cKO. (B) Representative RNAscope images showing *Scd1* expression in *Pdgfrα+* OPCs (green), BCAS1+ premyelinating, oligodendrocytes in control and OPC-Scd1 cKO corpus callosum at P7. Arrows indicate individual cells; note a strong reduction in the Scd1 expression in OPC-Scd1 cKO. (C-D) Representative images and quantification of EYFP and Olig2 labelled cells in control and OPC-Scd1 cKO brain sections at P10. (E-F) Representative images and quantification of *Pdgfrα+* OPCs in white matter and deep cortical regions from control and OPC-Scd1 cKO at P10. (G-H) *Plp1+* mature oligodendrocytes in the same regions as (E) show a significant reduction in the OPC-Scd1 cKO. Representative images are obtained from the highlighted regions shown in the cartoon insets. Scale bar: 25μm for images shown in (B) and 50μm for images shown in (C) and 100µm for images shown in (E and G). Data represent mean ± SEM, n=5-6 animals per group. ****p<0.0001, unpaired t-test.

Using an EYFP reporter line, we also confirmed that the *Pdgfrα-Cre*^*ER*^ line successfully targeted the oligodendroglial lineage cells (**Figure 1C**). Interestingly, we observed ∼20% reduction in the numbers of EYFP+Olig2+ cells in the *OPC-Scd1 cKO* mice compared to the control, suggesting a loss of oligodendroglial cell population (**Figure 1C-D**). Analysis of lineage specific markers revealed that while *Pdgfrα*+ OPC numbers remained unchanged (**Figure 1E-F**), we found a ∼50% reduction in *Plp1*-expressing oligodendrocytes in forebrain white matter regions of *OPC-Scd1 cKO* animals compared to controls (**Figure 1G-H**). This reduction in mature oligodendrocyte numbers without affecting the OPC pool indicates that *Scd1* is specifically required for the OPC differentiation. These findings raised the question of whether this differentiation defect would translate into myelination deficits.

### 3.2 Developmental Myelination Deficits in OPC-Scd1 cKO

Given the substantial reduction in *Plp1*+ oligodendrocytes, we next examined whether these cellular changes resulted in myelination deficits using markers for actively myelinating oligodendrocytes. Immunohistochemical analysis of Olig2 and CC1 markers revealed a modest but significant reduction in total Olig2+ oligodendrocyte lineage cells in *OPC-Scd1 cKO* animals at P10 (**Figure 2A-B**). More strikingly, the percentage of Olig2+ cells that were also CC1+ showed a ∼40% reduction in *OPC-Scd1 cKO* animals compared to controls (**Figure 2B**). This marked reduction in CC1+ mature oligodendrocytes confirmed that Scd1 deletion impairs the transition to active myelination. Consistent with the decrease in mature oligodendrocyte numbers, MBP and NF immunostaining also revealed reduced axonal myelination in both corpus callosum and striatum of *OPC-Scd1 cKO* animals (**Figure 2C-D**). The reduction in mature myelinating oligodendrocytes at P10 occurs during a critical CNS myelination window, which suggests that the desaturase activity of Scd1 to synthesize monounsaturated fatty acid is particularly important during the process of myelin sheath formation.

**Figure 2.**
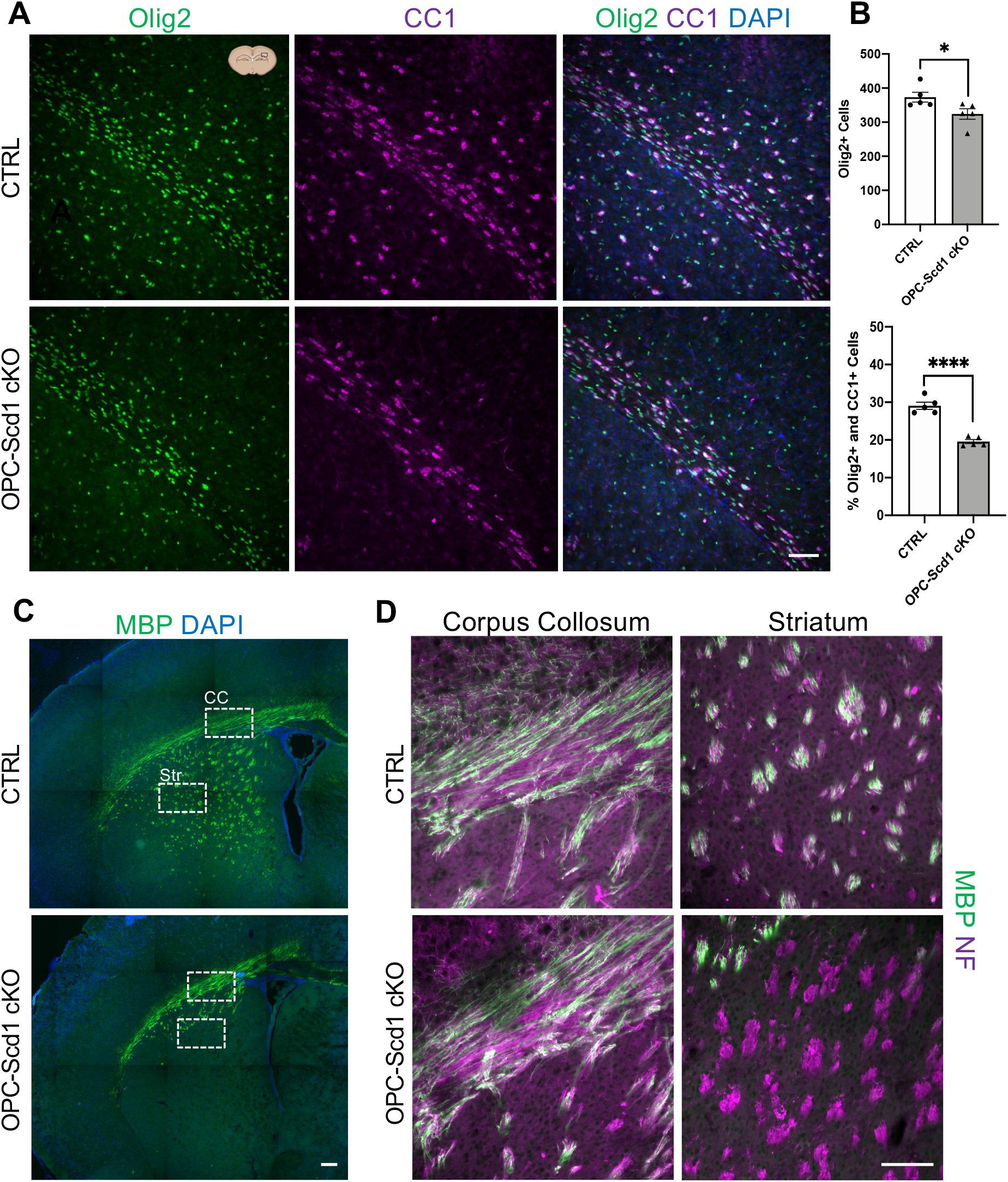
Developmental myelination deficits in OPC-Scd1 cKO mice. (A) Representative images of Olig2+ and CC1+ cells in white matter regions from control and OPC-Scd1 cKO mice at P10. Cartoon insets highlight the regions the images were obtained from. (B) Quantification of total Olig2+ cells and percentage of Olig2+CC1+ cells shows a significant reduction in mature oligodendrocytes numbers in OPC-Scd1 cKO at P10. (C) MBP immunostaining on coronal brain sections; highlighted regions are corpus callosum (CC) and striatum (Str) regions. (D) High magnification images of MBP and neurofilament (NF) immunostaining from the corpus callosum and striatum regions from (C) show axonal hypomyelination within the deep cortical and striatal regions. Scale bar: 50µm for images shown in (A) and 100μm for (C-D). Data represent mean ± SEM, n=5 animals per group. *p<0.05, ****p<0.0001, unpaired t-test.

### 3.3 RNA Sequencing and Fatty Acid Profiling Confirm Hypomyelination in OPC-Scd1 cKO

To comprehensively assess the molecular consequences of Scd1 deletion, we performed RNA sequencing on optic nerves from P10 control and OPC-Scd1 cKO animals (**supplementary table 1**). Gene ontology analysis revealed significant enrichment of myelination and oligodendrocyte development pathways among downregulated genes (**Figure 3A)**. Gene set enrichment analysis confirmed significant negative enrichment for oligodendrocyte markers (NES: -2.28, FDR: 0.0) and genes associated with myelin sheath formation (NES: -1.44, FDR: 0.004) (**Figure 3B**). Individual myelin-related genes showed consistent downregulation in *OPC-Scd1 cKO* samples (**Figure 3C**).

**Figure 3.**
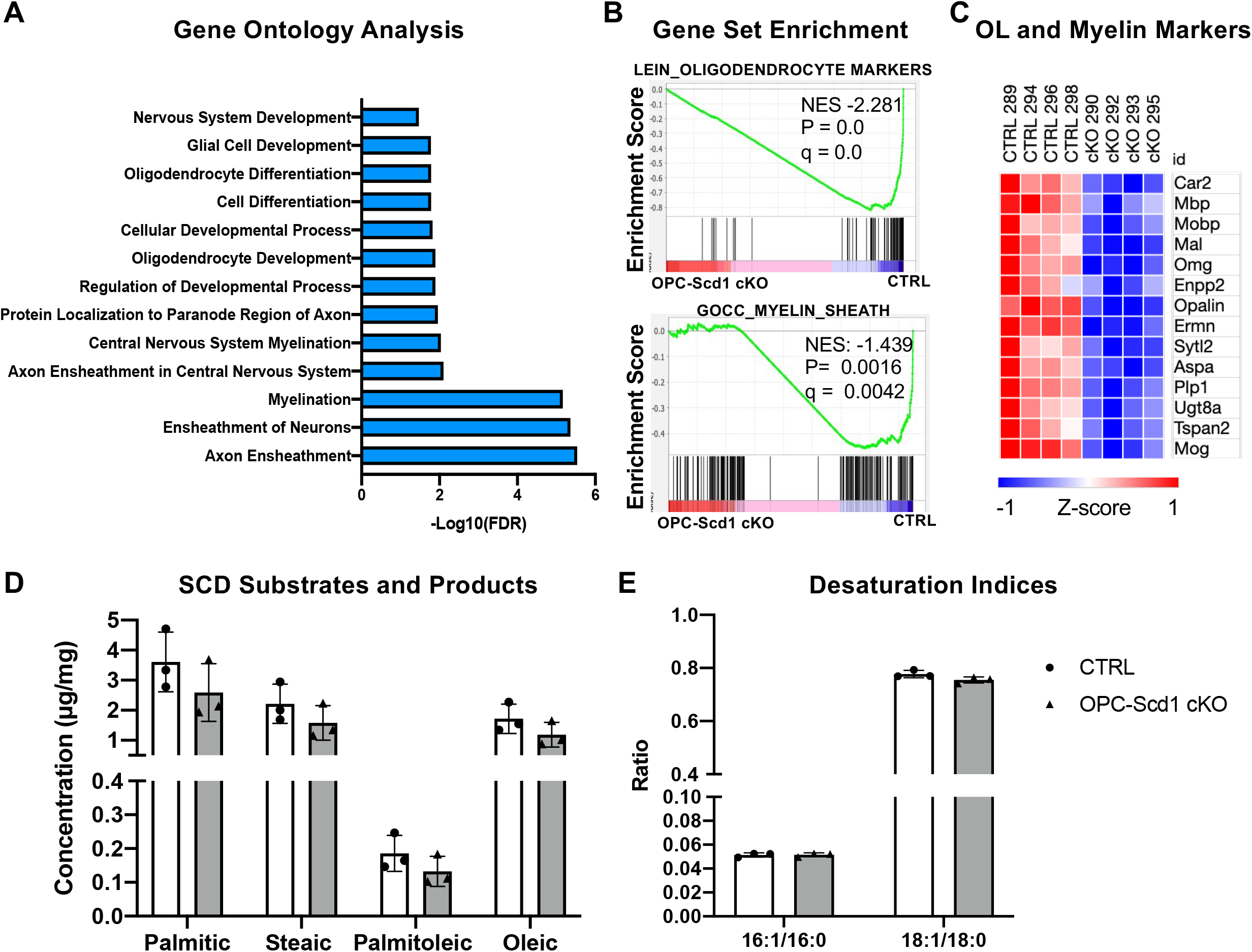
RNA sequencing and fatty acid profiling confirm hypomyelination in OPC-Scd1 cKO. (A) Gene ontology analysis of differentially expressed genes showing enrichment for myelination-related processes. (B) Gene set enrichment analysis (GSEA) plots for oligodendrocyte markers and myelin sheath gene sets with normalized enrichment scores (NES) and FDR q-values. (C) Heatmap of oligodendrocyte and myelin marker gene expression. (D) Fatty acid concentrations for Scd1 substrates (palmitic, stearic) and products (palmitoleic, oleic) in white matter isolates (corpus callosum and optic nerve tissue) from P10 control and OPC-Scd1 cKO mice. (E) Desaturation indices (16:1/16:0 and 18:1/18:0 ratios) calculated from the fatty acid concentrations in control and OPC-Scd1 cKO white matter isolates. Data represent mean ± SEM, n=4 for RNA-seq and n=3 animals per group for fatty acid analysis.

To complement the transcriptional analysis, we also performed fatty acid profiling of white matter tissue. For this, we isolated optic nerve and corpus collosum tissue from P10 control and *OPC-Scd1 cKO* mice. Consistent with the hypomyelination phenotype, we observed a uniform ∼30% reduction in all fatty acid species that were analyzed (**supplementary table 2**), including both Scd1 substrates (palmitic, stearic) and products (palmitoleic, oleic) (**Figure 3D**). Desaturation indices (16:1/16:0 and 18:1/18:0) remained unchanged in bulk tissue analysis (**Figure 3E**). Given that this analysis was performed on whole tissue rather than isolated oligodendrocytes, these findings likely reflect the tissue-level consequence of reduced myelin content rather than revealing cell-autonomous metabolic alterations.

### 3.4 Oligodendrocyte Maturation Delays Persist into Adolescence

To determine whether the developmental defects observed during development (P10) were transient or persistent, we extended our analysis to fourth postnatal week, which represents the adolescent period when myelination is more advanced but still ongoing in many brain regions. At this timepoint, *Pdgfrα*+ OPC numbers remained similar between control and knockout animals (**Figure 4A-B**), confirming that the OPC pool was maintained. Although *Plp1*+ oligodendrocyte numbers showed a trend toward reduction, this did not reach statistical significance (**Figure 4C-D**). Becausee *Plp1* mRNA is also expressed in newly differentiated oligodendrocytes, it is possible that these counts are not exclusively capturing mature population. To further assess oligodendroglial maturation, we performed IHC analysis using Olig2 and CC1 markers. We observed that the total Olig2+ cells showed no significant reduction (**Figure 4E-F**). However, the proportion of Olig2+CC1+ mature oligodendrocytes remained significantly reduced by ∼25% in *OPC-Scd1 cKO* animals (**Figure 4E-F**). Black Gold staining further confirmed modest but significant myelin reductions in both deep cortical and white matter regions (**Figure 4G-H**). The persistence of reduced CC1+ mature oligodendrocytes, despite recovery of total oligodendrocyte numbers, suggests that while some compensatory differentiation occurred, the timing and efficiency of maturation remained impaired. These findings prompted us to examine whether the cellular deficits translated into ultrastructural myelin abnormalities.

**Figure 4.**
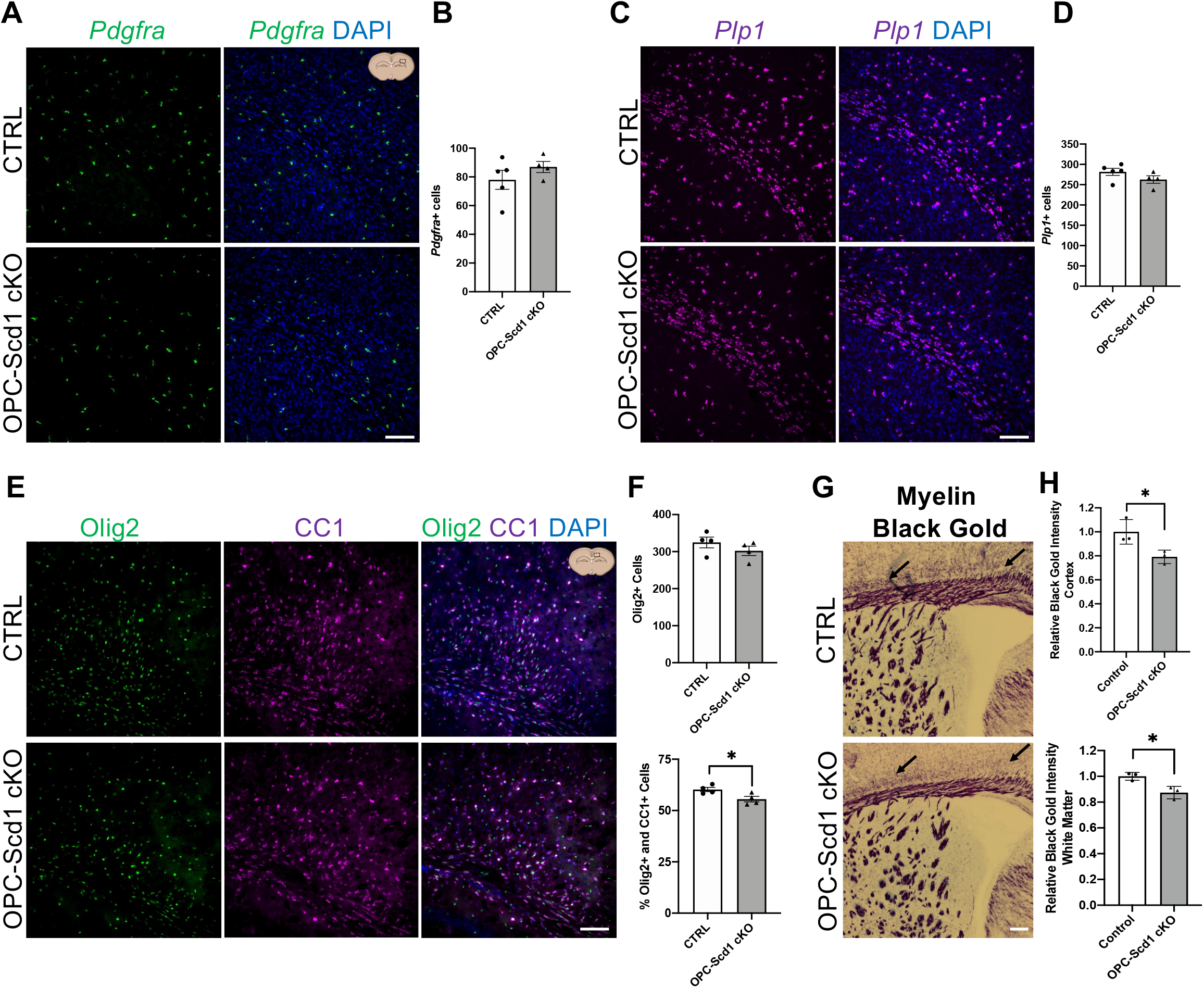
Oligodendrocyte maturation delays persist into adolescence. (A-B) Representative images and quantification of *Pdgfrα+* OPCs at P28. (C-D) *Plp1+* mature oligodendrocytes in P28 control and OPC-Scd1 cKO white matter and their quantification. (E-F) Olig2 and CC1 labeling shows showing persistent reduction in mature oligodendrocyte numbers at P28 in OPC-Scd1 cKO. (G-H) Staining and quantification of black gold in coronal brain sections show myelin reduction in deep cortical and white matter regions at P28 in OPC-Scd1 cKO (highlighted by arrows). Representative images are obtained from the highlighted regions shown in the cartoon insets. Scale bar: 100µm for images show in (A), (C) and (E) and 400µm for images shown in (G). Data represent mean ± SEM, n=4-5 animals per group. *p<0.05, unpaired t-test.

### 3.5 Axonal Myelination is Reduced in OPC-Scd1 cKO Optic Nerves

To assess the functional consequences of persistent oligodendrocyte maturation defects on myelin ultrastructure, we performed electron microscopy analysis of optic nerves isolated from P28 animals. The optic nerve provides a homogeneous population of myelinated axons ideal for quantitative analysis. G-ratio analysis on the myelinated axons revealed a modest but statistically significant increase from 0.72 to 0.75 in *OPC-Scd1 cKO* animals (**Figure 5A-C**). This shift indicated thinner myelin sheaths across the full range of axon diameters (**Figure 5B**), affecting both small and large caliber axons equally. Interestingly, we did not observe any significant abnormalities or structural differences in myelin wrapping around the axons between control and *OPC-Scd1 cKO* animals. Given the role of Scd1 in synthesis of monounsaturated fatty acids, which are known to play a role in membrane fluidity, our results showing myelin ultrastructure were surprising. Nonetheless, these findings were consistent with the cellular and molecular phenotypes observed and also raised the question of whether Scd1 would also be required for myelin maintenance in mature oligodendrocytes.

**Figure 5.**
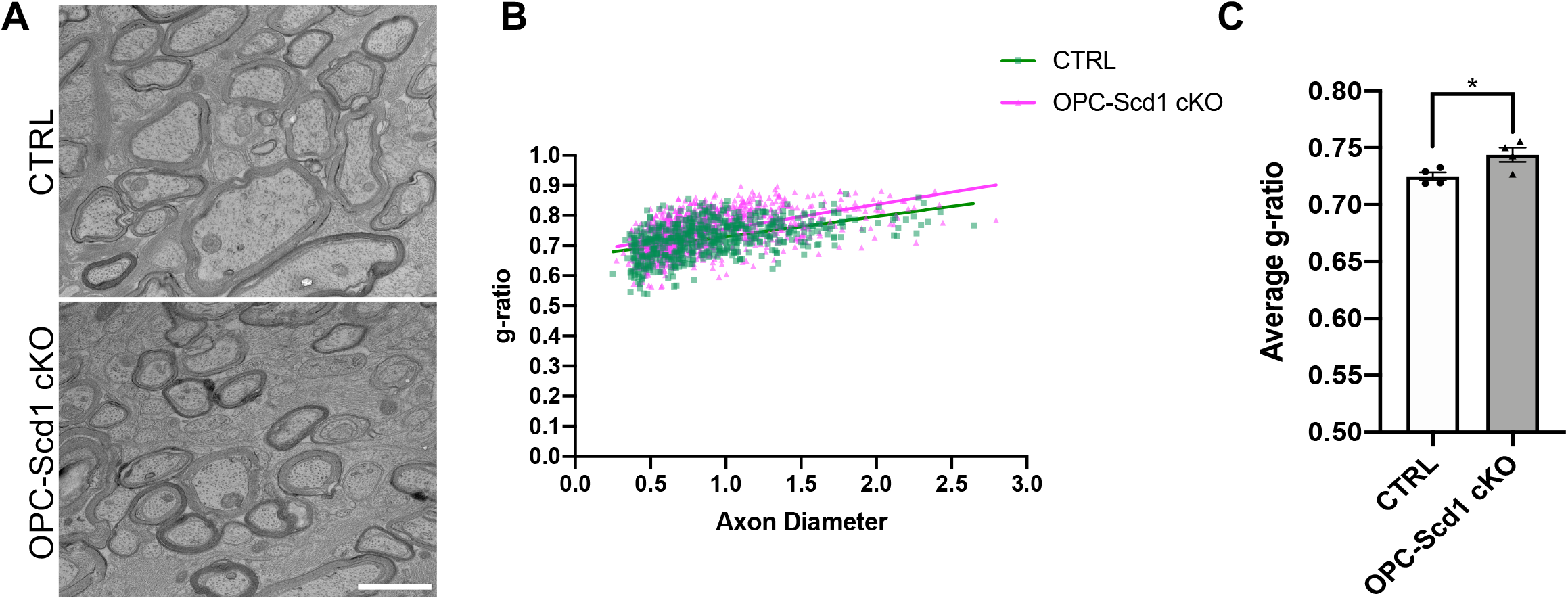
Optic nerve hypomyelination in adolescent OPC-Scd1 cKO mice. (A) Representative electron micrographs of P28 optic nerve ultrathin cross-sections. (B) Analysis of axonal myelination and scatter plot as a function of axon diameter showing increased g-ratios (thinner myelin) across all axon sizes. (C) Average g-ratio quantification. Data represent mean ± SEM, with 600-800 axons analyzed from n=4 animals/group. Scale bar: 1μm. *p<0.05, unpaired t-test.

### 3.6 Scd1 is Dispensable for Mature Myelin Maintenance

Given that Scd1 expression is highest in mature oligodendrocytes, we asked if it plays a functional role in either their survival or myelin maintenance. To determine this, we generated a conditional knockout model targeting Scd1 deletion specifically in mature oligodendrocytes using the tamoxifen inducible *Plp1-Cre*^*ER*^ line (referred to as *OL-Scd1 cKO*). Tamoxifen induction was first performed during early development at P13-P15, with analysis a month later at P45. Immunohistochemical analysis did not reveal any alterations in mature oligodendrocyte numbers and myelination (**Supplementary Figure 2**). Next, to understand whether Scd1 plays a role in myelin maintenance at later timepoints, we performed tamoxifen induction at P30 (**Figure 6A**) and analyzed the animals at P90 by electron microscopy. Remarkably, deletion of Scd1 in mature oligodendrocytes had no significant effect on myelin maintenance, as electron microscopic analysis revealed normal myelin ultrastructure in *OL-Scd1 cKO* (**Figure 6A**). The average g-ratio remained at ∼0.73 in both control and *OL-Scd1 cKO* animals (**Figure 6B**). In line with these results, we also did not observe any detectable changes in myelination by luxol fast blue and myelin black-gold staining (**Supplementary Figure 3**).

**Figure 6.**
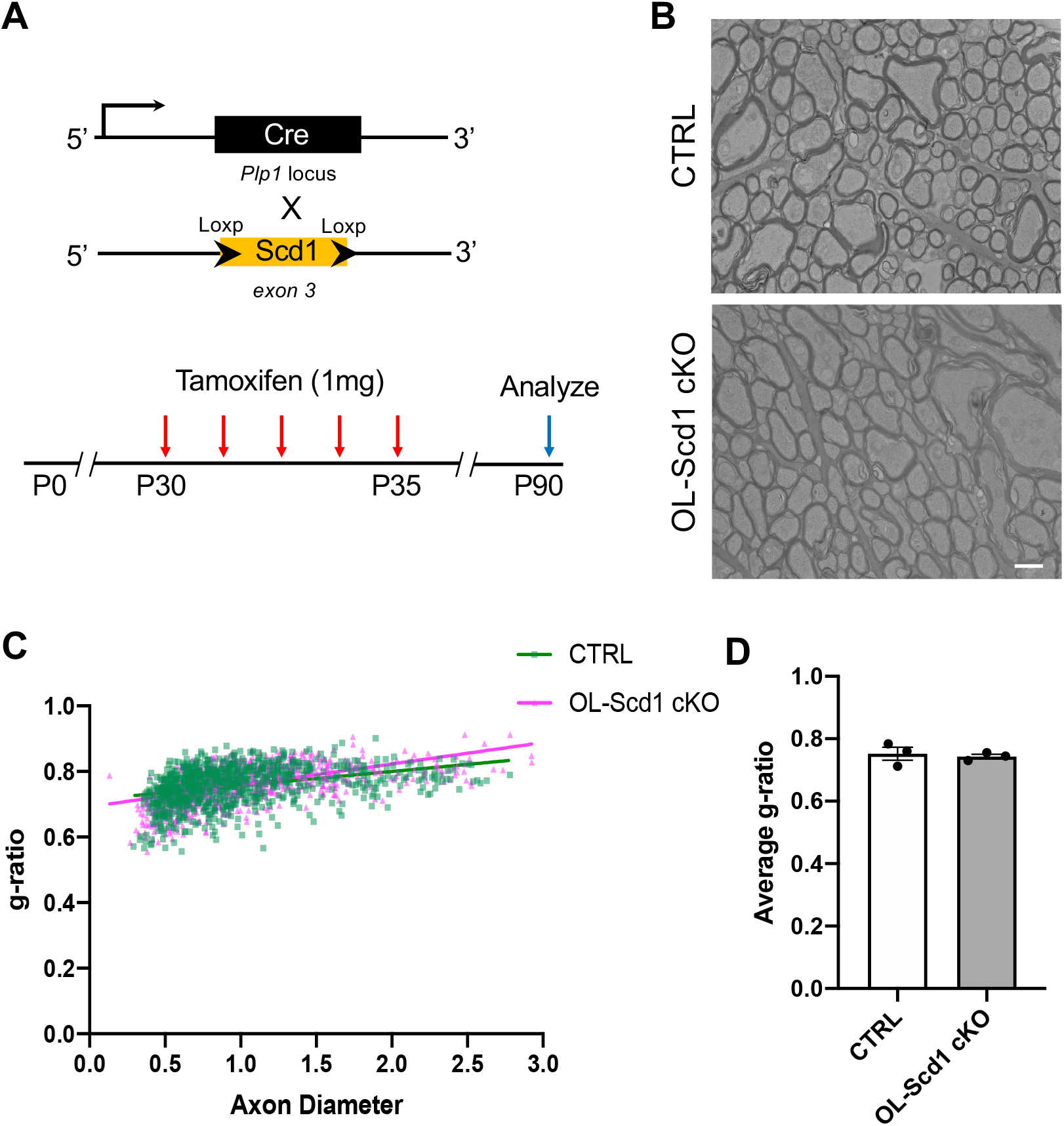
Scd1 is dispensable for mature oligodendrocyte and myelin maintenance. (A) Experimental design for Plp1-Cre^ER^-mediated genetic ablation of Scd1 (targeting exon 3) in mature oligodendrocytes (referred to as OL-Scd1 cKO). Tamoxifen administration from P30-P35 and analysis at P90. (B) Representative electron micrographs showing normal myelin ultrastructure and axonal myelination in OL-Scd1 cKO. (C) G-ratio distribution and (D) average g-ratio showing no difference between control and OL-Scd1 cKO. Scale bar: 1μm Data represent mean ± SEM, n=3 animals per group with 800-900 axons analyzed from 3 animals/group.

This striking difference between developmental and maintenance requirements indicated that Scd1 function is temporally restricted to the initial phases of myelination. The dispensability of Scd1 in myelin maintenance suggests metabolic requirements of oligodendrocytes change between the developmental periods and the maintenance phase. It also remains to be understood whether Scd2 activity is more critical for mature oligodendrocyte survival and myelin maintenance.

## Discussion

Our findings demonstrate that Scd1 plays a role in oligodendrocyte precursor cell differentiation and developmental myelination, while being dispensable for myelin maintenance in mature oligodendrocytes. This lineage and temporal specificity highlight the metabolic requirements of oligodendrocytes during different phases of myelination. The role of Scd1 during oligodendrocyte development is likely related to the massive myelin membrane expansion and biophysical properties that the monounsaturated fatty acids confer to myelin. During development oligodendrocytes generate lipid-rich myelin membrane required for axons of various calibers, making them one of the most metabolically active cell types within the CNS (Chrast et al., 2011; Montani, 2021, 2021; Naffaa et al., 2022; Saher et al., 2005; Young et al., 2013). The conversion of saturated fatty acids to monounsaturated fatty acids by Scd enzymes is crucial for maintaining appropriate membrane fluidity and facilitating axonal myelin wrapping (Schmitt et al., 2015; Min et al., 2009). The dispensability of Scd1 for mature myelin maintenance suggests the other isoform, Scd2 could be playing a compensatory role or is the sole desaturase within the mature oligodendrocyte population. Indeed, *Scd2* is expressed at much higher levels compared to *Scd1* within the oligodendroglial lineage cells, but whether *Scd1* prefers certain fatty acid substrates during early development remains to be explored.

The developmental delays in OPC maturation we observed showed partial recovery over time. Electron microscopy analysis also indicated thinner myelin sheaths but no structural defects in *OPC-Scd1 cKO* white matter tracts. The slow recovery of myelination deficits into adolescence (P28) indicates that the metabolic requirements for proper oligodendrocyte development are potentially compensated by Scd2 activity. This could suggest a potential metabolic redundancy, where Scd1 loss can be effectively compensated by Scd2 or other glial cell Scd1 mediated fatty acid and lipid transfer. The dispensability of Scd1 for myelin maintenance suggests that the metabolic demands for maintaining myelin differ from those required for initial myelin synthesis or a potential role for Scd2 in this process.

The molecular mechanisms by which Scd1 regulates oligodendrocyte differentiation require further investigation. A limitation in our study is that both RNA sequencing and fatty acid analysis were performed on bulk tissue rather than isolated oligodendrocytes. Our RNA sequencing analysis captures the tissue-level consequences of hypomyelination, and specific enrichment of myelination and oligodendrocyte development pathways in our gene ontology analysis, rather than broad metabolic disruption. This supports a targeted role for Scd1 in differentiation rather than general metabolic failure. Similarly, the uniform reduction in all fatty acid species coupled with unchanged desaturation indices likely reflects the fewer mature oligodendrocytes producing myelin, rather than revealing the metabolic state of individual cells. To understand the cell-intrinsic metabolic alterations caused by loss of Scd1, future studies on purified oligodendrocyte populations at different developmental stages would be required. Such analyses could reveal whether individual OPCs accumulate saturated fatty acid substrates, whether Scd2 compensates successfully in differentiating cells, or if specific lipid species critical for developmental differentiation are selectively affected.

Stearoyl-CoA desaturases are known regulators of Wnt ligand processing (Rios-Esteves & Resh, 2013) and given the established role of Wnt signaling in oligodendroglial maturation and white matter development (Hammond et al., 2015; S. P. Fancy et al., 2009; S. P. J. Fancy et al., 2011; Chavali et al., 2020; S. P. J. Fancy et al., 2014; Guo et al., 2015), investigating potential non-canonical functions of Scd1 could also provide additional mechanistic insights. In conclusion, our study demonstrates that Scd1 contributes to efficient oligodendrocyte precursor cell differentiation, with its loss resulting in a developmental myelination deficit that may be modulated by compensatory mechanisms. These findings reveal a temporal specificity in metabolic enzyme requirements within oligodendrocytes and further highlight the importance of fatty acid and lipid metabolism in their development. The differential requirement for Scd1 between developmental and maintenance phases also highlights the need to consider the developmental stage of target cells when developing therapeutic strategies to promote myelination in pathological contexts.

## Acknowledgments and Funding

MC acknowledges funding from NIH-R00NS117804, R00NS117804-S1 and First Tech Federal Credit Union Pediatrics Research Innovation Endowment.

