## Supplementary Figures for "Temporal Requirement for Stearoyl-CoA Desaturase 1 in Oligodendrocyte Development but Not Myelin Maintenance"

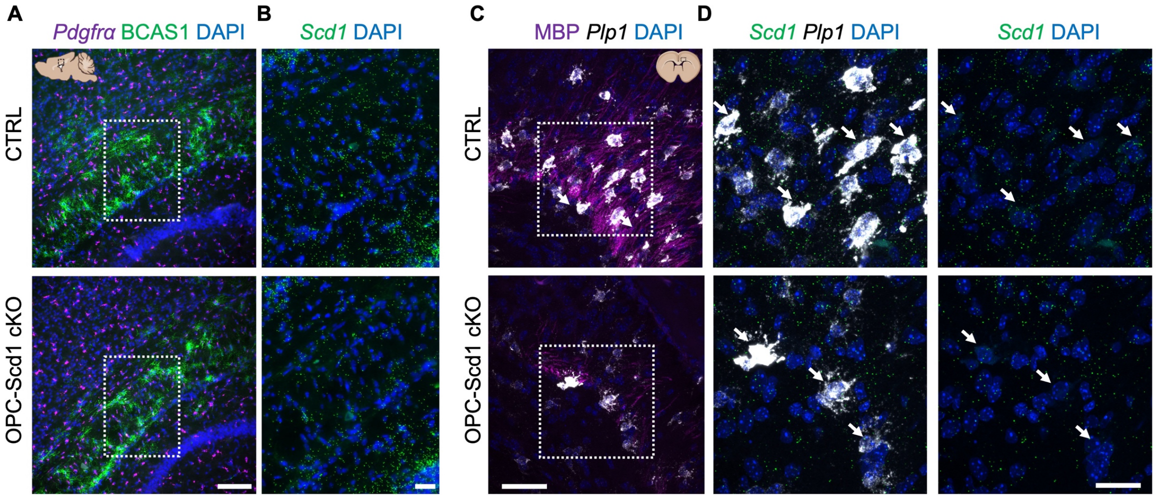


**Supplementary Figure 1. Scd1 Expression in White Matter** (A-B) Representative RNAscope images showing *Scd1* expression in P7 white matter along with *Pdgfrα*+ and BCAS1+ labelling of OPCs and premyelinating oligodendrocytes; note a reduction in the *Scd1* transcripts in OPC-Scd1 cKO white matter. (C-D) *Scd1* expression in *Plp1*+ oligodendrocytes in control and OPC-Scd1 cKO corpus callosum at P10. Arrows indicate individual *Plp1*+ cells in images shown in the right panel. Scale bar: 100μm for images shown in (A), 25μm in (B), 50μm in (C) and 25μm in (D).

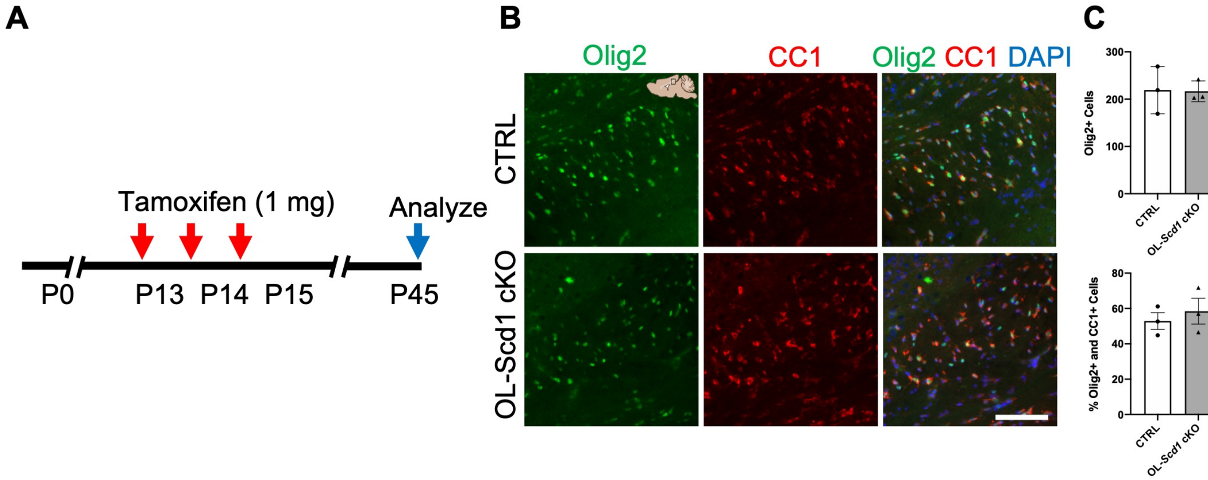


**Supplementary Figure 2. Scd1 is Dispensable for Mature Oligodendrocyte Maintenance** (A) Tamoxifen administration timeline in OL-Scd1 cKO from P13-P15 and analysis at P45. (B-C) Representative images of Olig2+ and CC1+ cells in the corpus collosum at P45 in control and OPC-Scd1 cKO show no significant differences numbers of mature oligodendrocytes Scale bar: 100μm. Data represent mean ± SEM, n=3 animals per group.

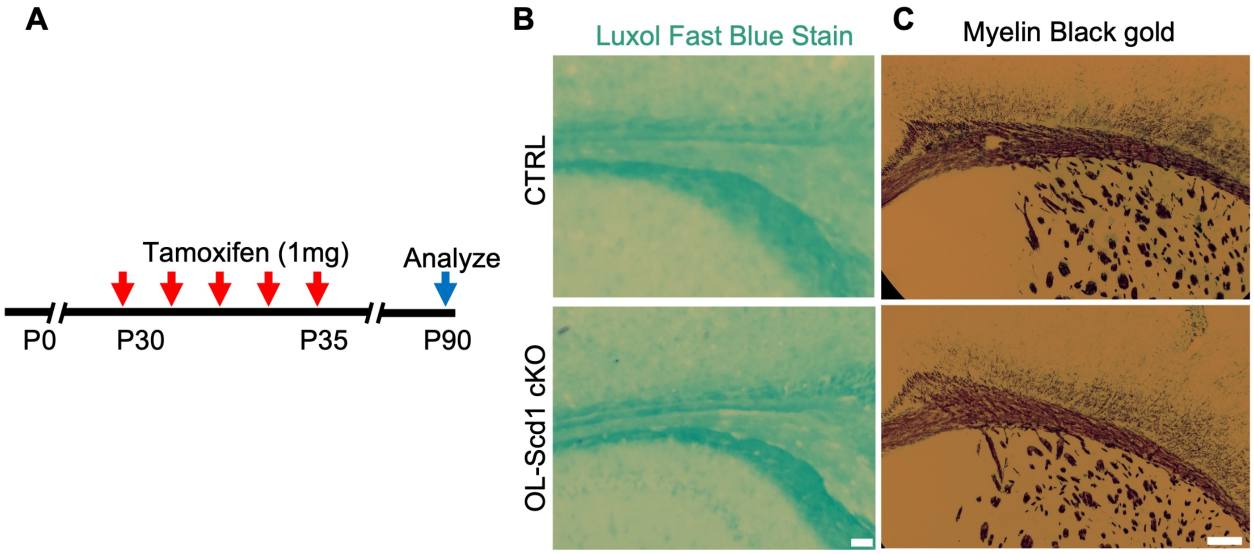


**Supplementary Figure 3. Oligodendrocyte maturation delays persist into adolescence.** (A) Tamoxifen administration timeline in OL-Scd1 cKO from P30-P35 and analysis at P90. (B-C) Representative images of luxol fast blue (B) and black-gold (C) staining show no alterations in the myelin levels in OL-Scd1 cKO. Scale bar: 100μm for images show in (B-C).
